# A molecular atlas of proximal airway identifies subsets of known airway cell types revealing details of the unique molecular pathogenesis of Cystic Fibrosis

**DOI:** 10.1101/2020.05.01.072876

**Authors:** Gianni Carraro, Justin Langerman, Shan Sabri, Zareeb Lorenzana, Arunima Purkayastha, Bindu Konda, Cody J. Aros, Ben A. Calvert, Aleks Szymaniak, Emily Wilson, Michael Mulligan, Priyanka Bhatt, Preethi Vijayaraj, Changfu Yao, David W. Shia, Edo Israely, Tammy M. Rickabaugh, Martin Mense, Scott H. Randell, Eszter K. Vladar, Amy L. Ryan, Kathrin Plath, John Mahoney, Barry R. Stripp, Brigitte N. Gomperts

## Abstract

Cystic fibrosis (CF) is a lethal autosomal recessive disorder that afflicts in excess of 70,000 people globally. People with CF experience multi-organ dysfunction resulting from aberrant electrolyte transport across polarized epithelia due to mutations in the cystic fibrosis transmembrane conductance regulator (*CFTR*) gene. CF-related lung disease is by far the most significant determinant of morbidity and mortality. In this study we report results from a multi-institute consortium in which single cell transcriptomics were applied to define disease-related changes to the proximal airway of CF donors (n=19) undergoing transplantation for end-stage lung disease compared to the proximal airway of previously healthy lung donors (n=19). We found that all major airway epithelial cell types were conserved between control and CF donors. Disease-dependent differences were observed, including an overabundance of epithelial cells transitioning to specialized ciliated and secretory cell subtypes coupled with an unexpected decrease in cycling basal cells. This study developed a molecular atlas of the proximal airway epithelium that will provide insights for the development of new targeted therapies for CF airway disease.

## Transcriptional analysis of single cells from control and CF airways

There is a great deal of interest in defining human bronchial epithelial (hBE) cell subtypes in Cystic Fibrosis (CF) airways as a means to develop gene therapeutic strategies to effect long-term correction of *CFTR* function. To address this knowledge gap we sought to produce single cell reference atlases of proximal airway epithelium isolated from lung tissue from donors with no evidence of chronic lung disease (CO, considered control for these experiments; n=19) compared to explant tissue from patients undergoing transplantation for end-stage CF lung disease (CF, n=19) (Supp Table1). Isolation of single cells from proximal airways was performed at three different institutions (Fig 1a), using similar but distinct methodologies (Fig 1b & Materials and Methods). After initial quality control and filtering, datasets from the three institutions were integrated for subsequent analyses. Data were visualized through UMAP dimensional reduction. The distribution of cells from each institution on UMAP projections showed homogeneous data integration (Supp Fig 1a, b). While all datasets integrated well, expression of some genes, particularly those associated with metabolic state, did show differential expression according to institution (Supp Fig 1c-f). Accordingly, data that were reproducibly observed across datasets from each of the three institutions were highlighted in this study.

**Figure 1.**
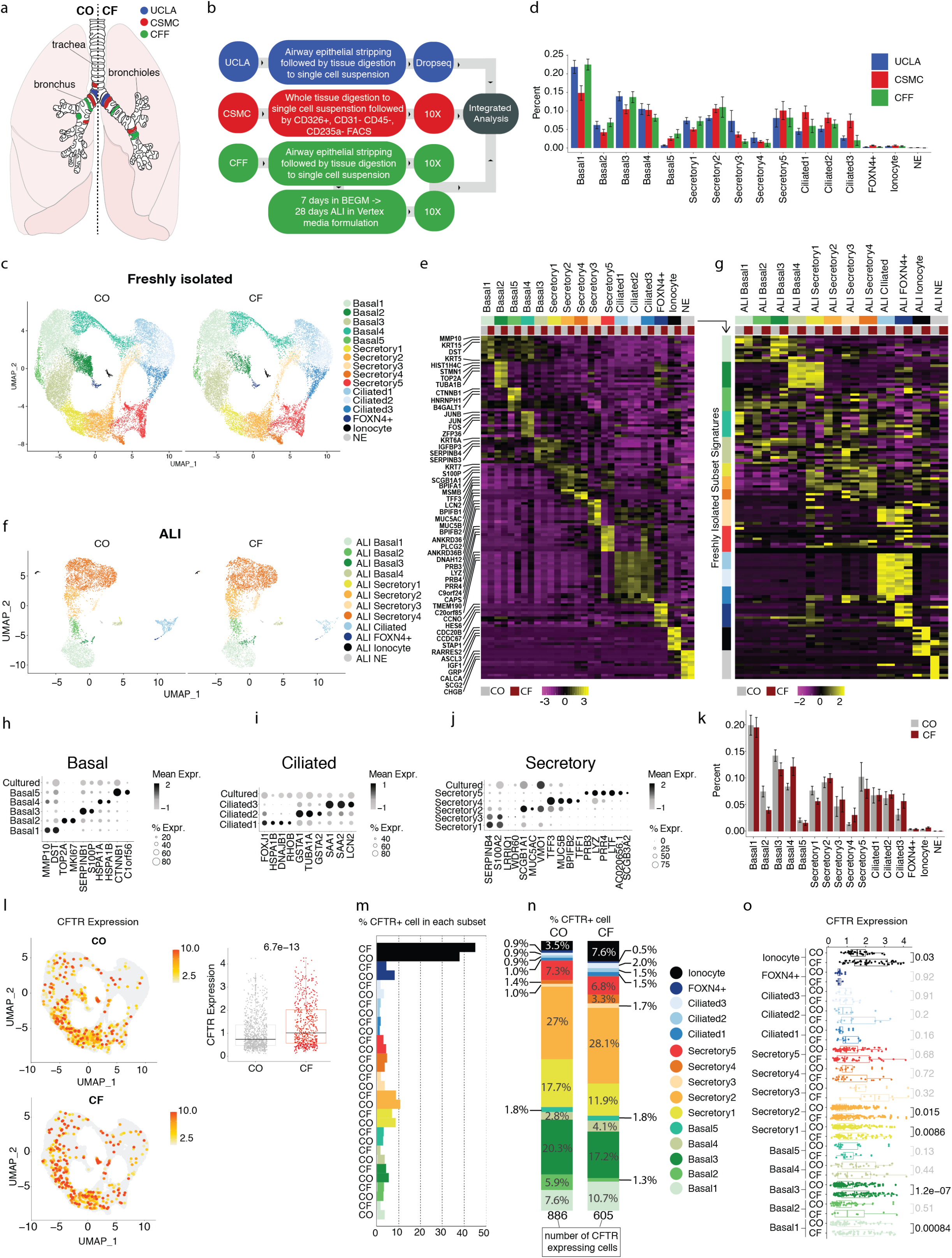
Single cell transcriptome atlas of the epithelium lining proximal airways of control donors and donors with end-stage CF lung disease. (a) Locations of cell procurement for single-cell RNA sequencing. (b) Methodology used for cell isolation by each institution. (c) Dimensional reduction of data generated from freshly isolated control and CF airway epithelium, visualized by UMAP, with cells colored by subtypes as shown in key. (d) Distribution of cell subtypes by institution. (e) Scaled expression of the top differentially expressed genes that inform specific cell subtypes, for k-groups of control and CF cells further separated by subtype, visualized by heatmap. (f) Dimensional reduction of data generated from air liquid interface cultures (ALI) derived from samples shown above. Cells are colored by ALI-specific subtypes, shown in key at right. (g) Heatmap of the scaled expression of the same fresh tissue subtype genes from (e), but shown for groups of ALI-control and CF cells split by subtype. (h-j) Comparison of subtype-specific gene expression among fresh tissue subtypes and cultured cells. (k) Distribution of the average proportion of cell subtypes per sample, comparing CO and CF cells (l-o) *CFTR* expression in subtype groups, key at right. (l) *CFTR* expression across all subtypes, shown on the UMAP projection and as a boxplot of CO/CF versus expression level (m) Proportion of *CFTR* expressing cells per each subtype. (n) Proportion of *CFTR*+ cells per cell subtype. (o) Boxplots showing the distribution of *CFTR* expression in all subtypes, for *CFTR*+ cells only, divided by CO and CF status. P values shown at right indicate the significance of distribution differences between CO and CF per subtype, bolded if p value < 0.05.

UMAP projections of data comparing cells from CO versus CF samples revealed a high degree of overlap (Fig 1c). Using the cell type gene signatures from Plasschaert et al^1^, we were able to identify all major human airway epithelial cell types including basal, secretory and ciliated, in addition to rare cell types including ionocytes, neuroendocrine (NE) and *FOXN4*+ cell populations (Supp Fig 1g,h). We then performed differentially expressed gene (DEG) analysis between clusters to discern cell subtypes with unique molecular characteristics. Among the three major cell types we were able to resolve 3 ciliated, 5 secretory, and 5 basal cell subtypes (Fig 1c, Supp Table2). These subtypes of each major airway epithelial cell type were found in similar proportions between CO versus CF samples and institutions (Fig 1d, Supp Fig 1i). We considered the functions conferred by differential genes to distinguish each cell subtype.

Basal cells were subdivided into five clusters (Basal1-5) (Fig 1e). The Basal1 cluster is characterized by high expression of canonical basal cell markers including tumor protein P63 (*TP63*) and the cytokeratins 5 and 15 (*KRT5* and *KRT15*) (Fig 1e, Supp Table2)^1^. Cells of the Basal2 cluster show enrichment for transcripts such as DNA Topoisomerase II Alpha (*TOP2A*) and the Marker of Proliferation Ki-67 (*MKI67*) and have a transcriptomic signature indicative of proliferating basal cells (Fig 1e). The Basal3 cluster is enriched for transcripts of the serpin family, members of which are known to regulate protein folding associated with secretory cell maturation and may represent basal cells transitioning to a secretory phenotype^2^. The Basal4 cluster is characterized by the highest expression of the AP-1 family members JUN and FOS, and Basal5 uniquely expressed β-catenin (*CTNNB1*).

Secretory cells were partitioned into five specific subsets (Secretory1-5) that share defining gene signatures in CO and CF datasets (Fig 1e). The Secretory1 cluster includes cells characterized by high expression of Secretoglobin Family Member 1A1 (*SCGB1A1*) and various members of the Serpin family. Serpins regulate protein folding associated with maturation of secretory proteins^2^ and define cells undergoing maturation into secretory cell type with similarities to club cells of bronchiolar airways^3^. The Secretory2 cluster is composed of cells sharing expression of mucins *MUC5B* and *MUC5AC*, anterior gradient 2 (*AGR2*) and SAM-pointed domain–containing Ets-like factor (*SPDEF*), suggesting they are goblet cells^4^. Cells in the Secretory3 cluster lack expression of known canonical secretory cell markers and can be distinguished from other secretory cluster subsets by their expression of Dynein Axonemal Heavy Chain proteins (*DNAHs*), Ankyrin Repeat Domain proteins (*ANKRDs*), and the mucins *MUC16* and *MUC4*, suggesting that the Secretory3 cluster acts as progenitor cell for ciliated cell differentiation. The Secretory4 cluster is defined by expression of *MUC5B* and Trefoil Factor family domain peptides (*TFF1* and *TFF3*) and represents a subtype of mucous-like cells that is distinct from goblet cells^5^. The Secretory5 cluster contains a serous-like signature^5^, including expression of Lysozyme (*LYZ*), Proline-Rich Proteins (*PRBs*, and *PRRs*), and Lactoferrin (*LTF*), and may represent glandular cell types of submucosal glands (SMGs) or their surface airway epithelial counterparts (Supp Table2).

Ciliated cells were subdivided in to three clusters (Ciliated1-3) (Fig 1e), all sharing expression of the lineage marker / master regulator of ciliogenesis, Forkhead box protein J1 (*FOXJ1*)^6^. The Ciliated1 cluster contains the highest expression of markers of cilia pre-assembly^7^, including Sperm Associated Antigen 1 (*SPAG1*), Leucin Rich Repeat Containing 6 (*LRRC6*) and Dynein Axonemal Assembly Factor 1 (*DNAAF1*), whereas cells within the Ciliated2 cluster show the highest expression of markers of mature ciliated cells including *TUBA1A* and *TUBB4B*. The Ciliated3 cluster is characterized by the expression of Serum Amyloid A proteins (*SAA1* and *SAA2*), reflective of a pro-inflammatory state^8^, suggesting that this subset of ciliated cells is either responding to or regulating immune responses.

In light of the cellular heterogeneity observed among freshly isolated airway epithelial cells we sought to determine the extent to which this was recapitulated in commonly used culture models, most notably the primary human bronchial epithelial (hBE) cell air liquid interface (ALI) culture system. We performed single cell RNA sequencing on well differentiated ALI cultures^9^ generated from hBE cells matched to a subset of CO and CF donors used for analysis of freshly isolated cells. Previously identified cell types^10^ observed in fresh isolates (basal, secretory, ciliated, *FOXN4*+, ionocyte, and NE) were also observed in ALI cultures (Supp Fig 1j), for both CO and CF-derived samples (Supp Fig 1k). Based on gene expression differences in ALI cultures, we were able to further define subtypes of basal (ALI Basal1-4), secretory (ALI Secretory 1-4), and ciliated cells (ALI Ciliated 1) (Fig 1f). ALI Basal1, 2, and 4 showed overlapping marker gene expression with Basal1 (Canonical), Basal3 (Serpin-enriched), and Basal2 (proliferating) cells from freshly isolated tissue, respectively (compare Fig 1e and 1g, Supp Table2, S3). ALI Basal3 identified cells with high *KRT14* expression that lacked a counterpart basal cell cluster in the fresh tissue data sets (Fig 1e, g). ALI secretory and ciliated cell clusters lacked markers observed in the respective subtypes of the freshly isolated tissue (Fig 1e, g, Supp Table3). The comparison of gene expression profiles between cells from ALI cultures and fresh tissue confirm that while the major cell types are present in ALI cultures, significant differences are observed in subtype states (Fig 1h,i, j). We conclude that ALI cultures recapitulate major hBE cell types observed among freshly isolated airway epithelial cells, as a regenerative model, but the cultures do not fully recapitulate the heterogeneity of cell subtypes observed in native airways at steady state.

Despite differences across the donors, isolation techniques, and sequencing methods, we found all cell subtypes in fresh airway tissue were recapitulated in each CO and CF patient (Fig 1k, Supp Fig 1k). Interestingly, we observed an average proportionate depletion in CF samples of 46.8% less cells in the proliferative Basal2 subtype and of 26% in the club cell-like Secretory1 subtype compared to CO, while a 44.6% increase in the proportion of cells in the inflammatory Ciliated3 subtype was observed (Fig 1k). These changes were observed in all three institutions, showing that the rigorous subtyping of human bronchial epithelium allows the identification of reproducible differences in cell states between CF and CO airways.

We next used our molecular atlas to examine cystic fibrosis transmembrane regulator (*CFTR*) expression in different cell types in CO and CF airways. Recent studies have proposed ionocytes as specialized cells with high *CFTR* expression that may represent primary tractable targets for restoration of *CFTR* expression in CF^10,11^. We found that *CFTR* is expressed in a wide selection of cells, with overall higher expression in CF compared to CO (Fig 1l). While more than 30% of all ionocytes expressed *CFTR* in both CO and CF samples, the majority of *CFTR*-expressing cells were secretory cells, followed by basal cells, with ionocytes constituting a minor fraction (Fig 1m,n). Analysis of the proportional expression of *CFTR* among secretory and basal cells showed that secretory cells and not ionocytes are the major producer of *CFTR* in both CO and CF tissue (Supp Fig 1l). Secretory2 (goblet-like) cells and Basal3 (serpin-expressing) cells were the major contributors to *CFTR* expression among the identified cell subtypes (Fig 1n, Supp Fig 1l). The comparison of *CFTR* expression between CO and CF samples showed cell type-specific differences, with increases of expression in CF samples in the ionocyte, Secretory1 (Club-like), Secretory2 (Goblet-like), Basal1 (Canonical), and Basal3 (serpin-expressing) cell subtypes (Fig 1o). Overall, our analysis, while confirming the specialized role of ionocytes for *CFTR* expression, establishes that secretory cells are the main cells that express *CFTR* and that secretory and basal cells together contribute the vast majority to *CFTR* expression in the proximal airway epithelium. Therefore, both secretory and basal cells should be considered as plausible candidates for therapeutic restoration of *CFTR* expression in CF in addition to ionocytes.

## Secretory cells show a secretory signature with increased antimicrobial activity in CF donors

The secretory cells of the proximal airway epithelium play an important role in host defense by producing serous and mucous secretions that trap and clear microbial organisms. People with CF develop dehydrated mucus and problems with mucus clearance which result in chronic bacterial infections. We explored the gene expression differences of secretory cell subtypes between CO and CF samples to define transcriptional differences that may be associated with these issues.

In order to find differences in each secretory cell subtype, we applied DEG analysis to identify the most specifically expressed genes in a given secretory subtype in CO or CF donors, and selected gene expression changes that were cross validated between all three datasets (Fig 2a, Supp Table2). In the Secretory1 (Club-like) subtype, CF samples showed downregulation of members of the *S100* gene family^12^, which are important for tissue repair, differentiation and inflammation, suggesting possible repair defects in CF donors. In the Secretory2 (Goblet-like) subtype, we found that immune response genes such as *BPIFA1* and *BPIFB1*^13^ were upregulated in CF samples. The Secretory3 (*DNAHs* enriched) subtype shows CF-specific increased expression of specific dyneins (*DNAH5,11,12, DNAAF1*), transcripts that are usually associated with cilium assembly^14^. In the Secretory4 (mucous-like) subtype, Angiogenin (*ANG*) and *TFF1*, two molecules with a role in antimicrobial defense^15,16^, were upregulated in CF compared to CO samples. Secretory5 (serous-like) subtype showed fewer differences between CO and CF compared to other subtypes (Fig 2a).

**Figure 2.**
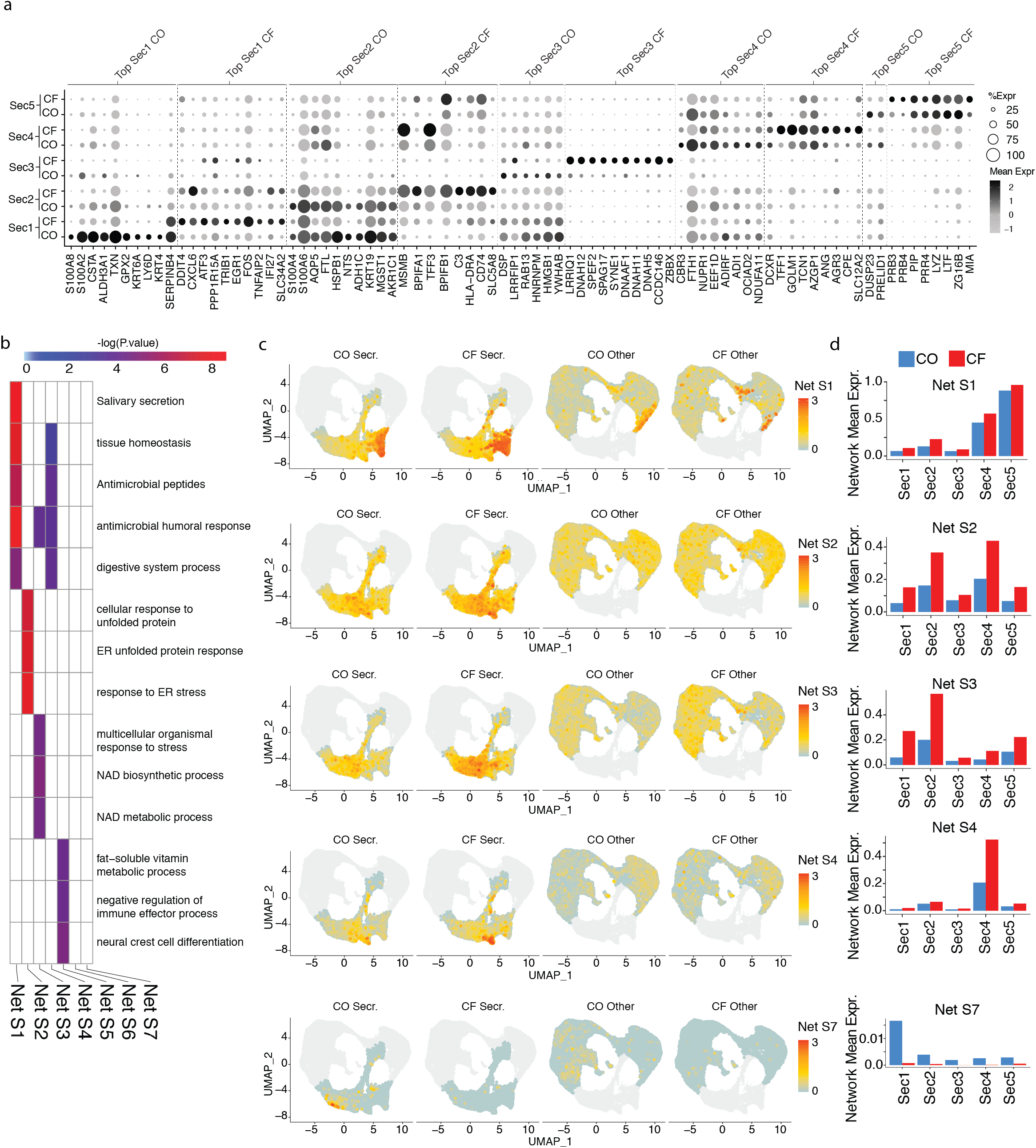
Expansion of secretory function, including mucus secretion and antimicrobial activity, in cystic fibrosis secretory cells. (a) Dot plot indicating the expression level and frequency of differentially expressed genes from each secretory subtype, across all subtypes in CO and CF cells. Genes are expressed higher in either CO or CF, as indicated by label at top. (b) For gene networks preferentially located in secretory cells, shown is a gene ontology heatmap of the top 3 associated terms for each network with the term enrichment −log(p-value) colored as displayed in key. Networks with no associated ontology terms are blank (Net S6/S7). (c) For each cell, the average mean expression of the genes in a given network is shown, visualized on a UMAP. Cells are split by Secretory or non-Secretory, and CO or CF classification (d) Bar plots showing the average expression of all genes in individual secretory networks per secretory subtype, in CO or CF cells.

To better understand the alterations to co-regulated sets of genes in CO compared to CF samples, we additionally applied an unbiased gene expression network discovery method that employs correlation between transcript levels to group genes to define co-expression networks that are most prominently expressed in secretory cells. We focused on seven of these networks (Net S1-S7) with statistical significance for differences in CO versus CF samples across each institution’s data (Fig 2b, Supp Fig 2, Supp Table4). Secretory networks 1-6 (Net S1-S6) are more highly expressed in CF vs CO secretory cells, in particular S2-S4, whereas S7 was lower in secretory cells from CF samples (Fig 2c, Supp Fig 2). Gene ontology analysis revealed an antimicrobial signature^17^ for S1 and S4, S2 was related to ER stress^18^ and S3 linked to metabolic processes (Fig 2b). The antimicrobial gene program S1 was most highly expressed in the Secretory4 and Secretory5 subgroups and expression of S4 was high specifically within the Secretory4 cluster (Fig 2c,d). This indicated that the Secretory5 (serous-like) and Secretory4 (mucous-like) cells in CF lungs have a highly specialized response in a particular subtype of cells and increased levels of antimicrobial activity in response to disease. Elevated ER-stress seen with S2 was more pronounced among Secretory4 and Secretory2 (goblet-like) cells (Fig 2c,d). S3 described a metabolic difference between Secretory2 (goblet-like) and Secretory1 (club-like) cells from CF versus CO samples (Fig 2c,d), indicating the surface airway epithelial secretory cells may be more exhausted. S5, marked by developmental ontology and containing the Wnt signaling gene *FRZB*, and S6, which contained Notch gene *HEY1*, were also elevated in CF samples (Supp Fig 2). Only one secretory network, S7, had a notable upregulation in CO compared to CF samples and marked a small group of cells expressing members of the *KLK* family, reported to be expressed in epithelial cells of the lung^19^, and implicated in regulation of airway inflammatory response (Fig 2c,d).

Overall, gene expression and network differences identified between CO and CF secretory cell subtypes demonstrate overactive mucosal secretion, humoral immunity, antimicrobial activity and stress-related organelle maintenance, suggesting an increase in secretory function in the CF airway epithelium.

## An expanded ciliated cell gene expression program reveals aberrant ciliogenesis and altered cellular lineages in CF airways

Multi-ciliated cells provide the necessary mechanical force for the directional clearance of contaminants trapped in the mucus layer for optimal airway homeostasis and host defense. Although CF-related defects in electrolyte transport across the epithelial lining and associated dehydration of airway surface liquid impair ciliary function and muco-ciliary transport, we know little about how ciliated cells respond to this perturbation. To comprehensively assess CF-related changes in ciliated cells, we compared their single cell transcriptomes between CO and CF patient samples. Ciliated cell gene expression is driven by a complex multi-ciliated cell specific gene expression network that turns on downstream of cell fate acquisition to generate the hundreds of structural and regulatory components of cilia^20^. In general, the temporal expression of specific ciliary genes reflects their role in sequential stages of ciliogenesis, and many transcripts are first strongly induced, then are eventually downregulated to a maintenance expression level^21^. Differential gene analysis revealed genes that were specific to either CO or CF cells in each of the ciliated subtypes, again reproducible between datasets from all three institutions (Fig 3a). The Ciliated1 subtype (cells undergoing ciliogenesis) showed higher expression for ciliogenesis transcripts such as Dynein Axonemal Heavy Chain 5 (*DNAH5*), Spectrin Repeat Containing Nuclear Envelope Protein 1 and 2 (*SYNE1* and *SYNE2*) in CF compared to CO tissue, suggesting an attempt to boost cilium biogenesis in lungs from CF donors. Cells of the Ciliated2 (mature ciliated) subtype, showed higher expression of Anterior Gradient 3 (*AGR3*) and *LRRC6* in CF samples, genes that play a role in ciliary beat frequency and motility^22,23^. CF cells of the Ciliated3 (cells involved in immune defense) subtype showed higher expression of Major Histocompatibility Complex, Class II, DP Alpha 1 and DR Beta 1 (*HLA-DPA1* and *HLA-DRB1*), genes that play an important role in the immune system in antigen presenting cells.

**Figure 3.**
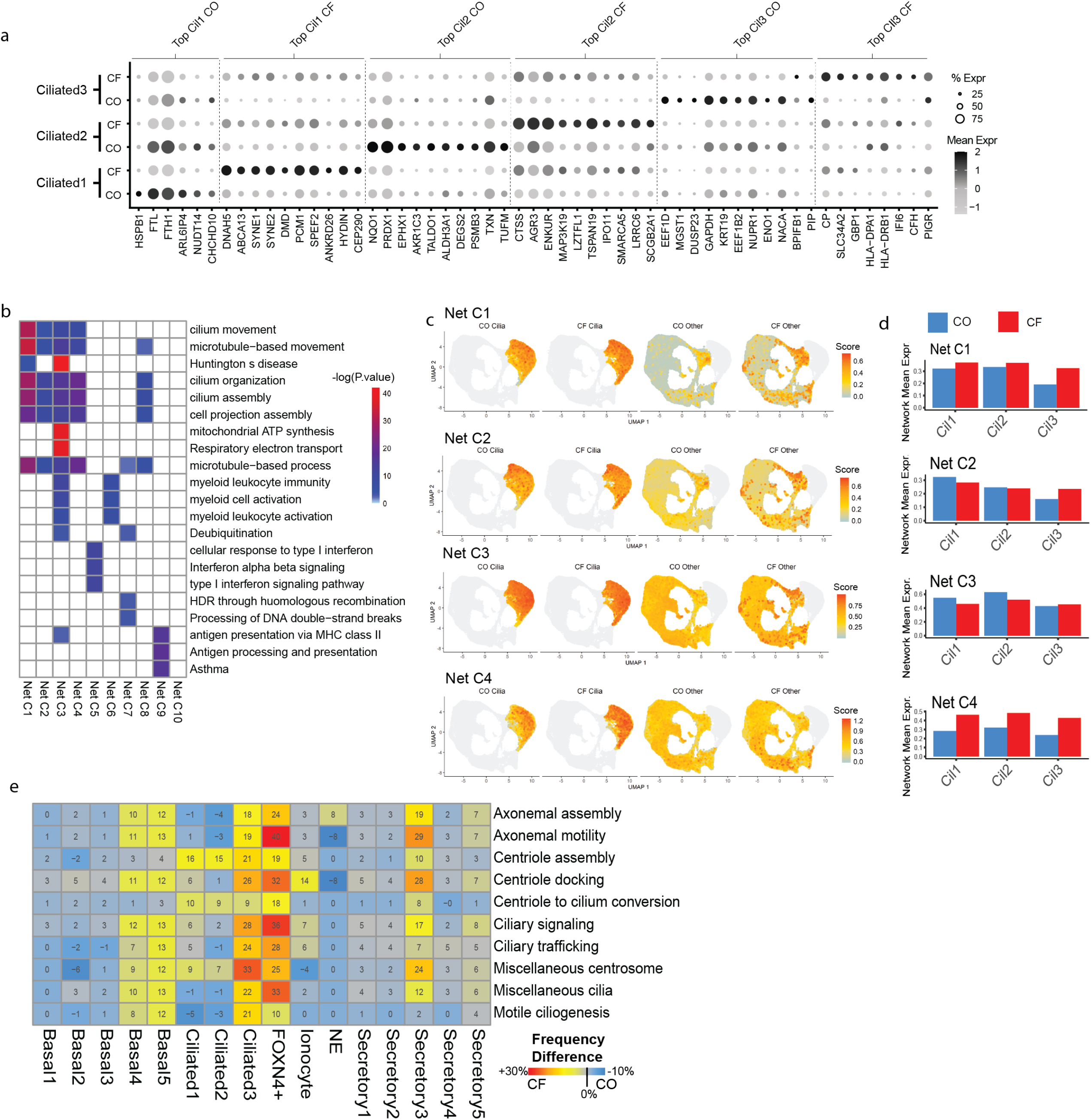
Cilia related gene expression is vastly expanded outside of the main cilia subgroups in CF. (a) Dot plot indicating the expression level and frequency of differentially expressed genes in each ciliated subtype, for CO or CF cells. (b) For gene networks preferentially expressed in ciliated cells, shown is a gene ontology heatmap of the top 3 associated terms for each network with the term enrichment −log(p-value) colored as displayed in key. (c) For each cell, the average mean expression of the genes in a given network is shown, visualized on a UMAP. Cells are split by Ciliated or non-Ciliated, and CO or CF classification (d) Bar plots showing the average expression of all genes in individual ciliated networks per ciliated subtype group, in CO or CF cells. (e) For distinct categories of genes related to cilia biogenesis, the expansion of cilia gene expression is shown by a heatmap indicating the proportional percent change in amount of cells in each subtype expressing each category above a threshold, towards CF(+%) versus CO(-%) cells. The percent change number between CF and CO samples is given in each heatmap cell and colored as indicated in key at right.

Through gene expression network discovery, we also defined ten expression networks that are highly expressed in ciliated cells (Fig 3b, Supp Fig 3). Despite each network having distinct genes, many networks showed enrichment of ontology terms related to ciliogenesis and cilium movement (Net C1-C4, C8; Fig 3b, Supp Fig 3, Supp Table4). Network C3 was associated with respiratory electron transport, C7 related to cellular repair and networks C3, C5, and C6 contained genes with immune functions (Supp Fig 3). The smaller networks C9 and C10 had no ontology but also contained immune and ciliary genes (Supp Fig 3). Interestingly, the Ciliated3 subtype consistently showed an increase in expression of all of these networks in CF compared to CO cells and often displayed the largest gene expression difference between these samples (Fig 3c,d; Supp Fig 3). We also found that the microtubule and ciliogenesis-related networks C1-C4 and C8 had higher expression among non-ciliated cells in CF compared to CO tissues (Fig 2b, c).

Given this specific and unexpected upregulation of various cilium-related genes in non-ciliated cells in CF samples, we wondered if particular cell subtypes were affected. To address this question, we interrogated a manually curated list based on published observations, containing a total of 10 categories and 491 genes, representing different phases of ciliogenesis (Fig 3e, Supp Fig 4). We then compared the proportion of cells from each hBE cell subtype that expressed a given ciliogenesis signature above a specific cutoff between CO and CF samples. For the Ciliated1 and Ciliated2 subtypes, we found that more cells expressed centriole-assembly and centriole-to cilium conversion gene signatures in CF than in CO samples suggesting a defect in ciliogenesis kinetics. All 10 ciliogenesis signatures were expressed by a higher proportion of CF cells in the “immune defense” Ciliated3 cell subtype, suggesting aberrant regulation of ciliogenesis in this subtype of ciliated cells in CF airways.

Among non-ciliated subtypes, Basal4, Basal5 and Secretory3 clusters had higher expression of nearly all categories of ciliogenesis signature genes in CF compared to CO samples, indicating a potential commitment toward the ciliated lineage in these CF non-ciliated cells (Fig 3e). Interestingly, *FOXN4*+ cells, previously reported to represent transitional *FOXJ1*+ cells undergoing multiciliogenesis^10^, were also found to express ciliogenesis signature genes at a higher level in CF compared to CO samples. Taken together, these data suggest that CF airways include an overabundance of cells transitioning to the ciliated cell phenotype compared to CO airways. We speculate that this may result in generation of more structurally and functionally aberrant ciliated cells and more immune defensive Ciliated3 cells in CF tissues. Furthermore, the epithelial lining of CF airways exhibits a more plastic and stressed phenotype consistent with known airway defects resulting from electrolyte and ASL imbalances in the CF airway.

## CF basal cells show depletion of metabolic stability and proliferation

Basal cells are considered to be the primary stem cells of the proximal airways that are capable of proliferation, long-term self-renewal, and differentiation to yield specialized luminal cell types^24,25^. Analysis of differentially expressed genes between basal cells of CO and CF samples revealed reproducible subtype-specific differences (Fig 4a). The CF Basal2 (Proliferating) cell subtype showed a general reduction of transcripts involved in cell division, whereas the CF Basal3 (Serpin-expressing) subtype showed lower expression of keratinization-associated genes^26,27^ including Cystatin A (*CSTA*) and Heat Shock Protein Family B (Small) Member 1 (*HSPB1*). The CF Basal4 (activated) subtype displayed increased expression of Fos and FosB Proto-Oncogene, AP-1 Transcription Factor Subunit (*FOS*, and *FOSB*), whereas the AP-1 complex companion transcription factors Jun and JunB (*JUN* and *JUNB*) were unchanged.

**Figure 4.**
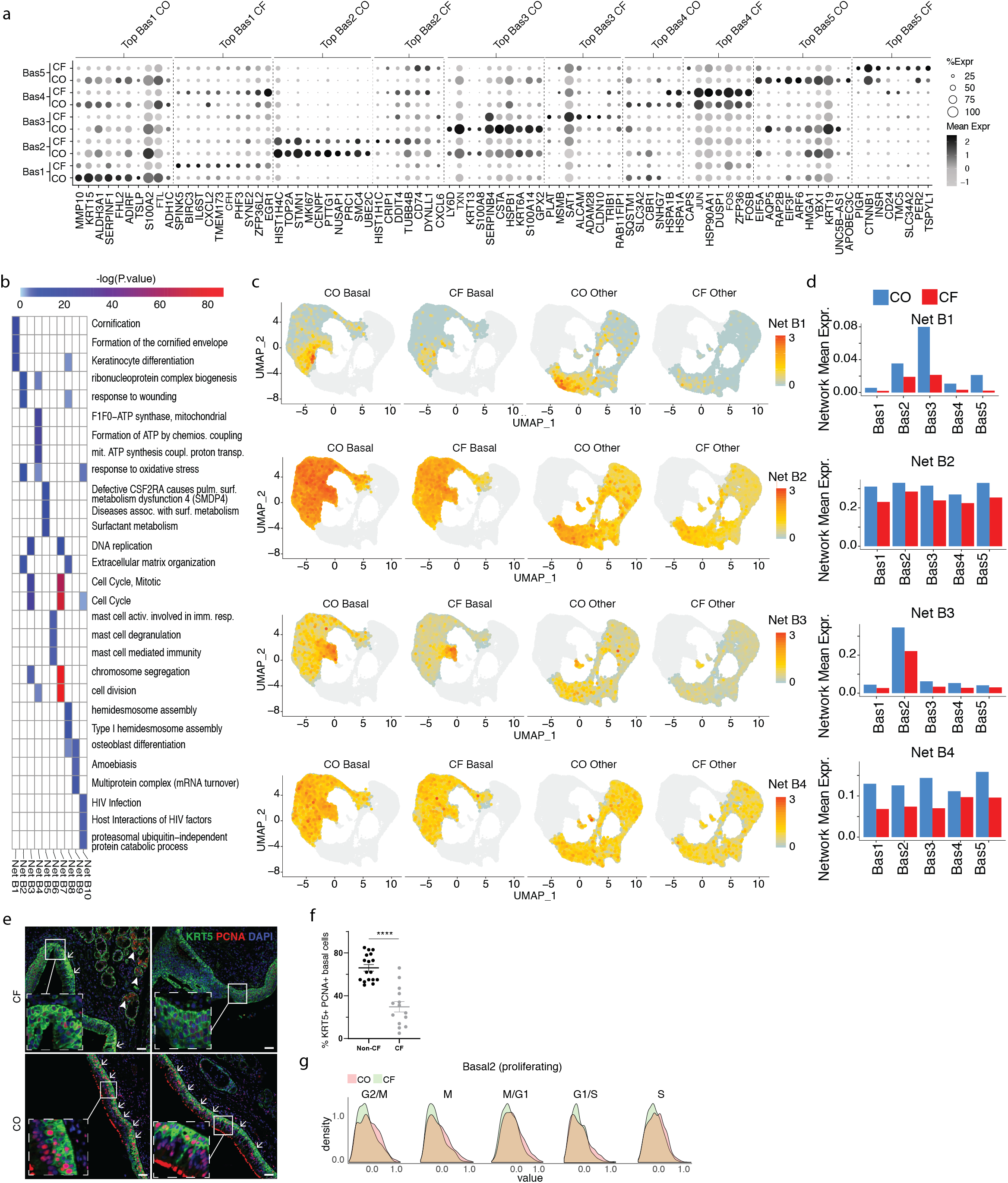
Depletion of metabolic stability, basal epithelial function, and cellular division is widespread in CF lung basal cells. (a) Dot plot indicating the expression level and frequency of differentially expressed genes in each basal subtype, for CO or CF. (b) For gene networks highly expressed in basal cells, shown is a gene ontology heatmap of the top 3 associated terms for each network with the term enrichment −log(p-value) colored as displayed in key. (c) For each cell, the average mean expression of the genes in a given network is shown, visualized on a UMAP. Cells are split by Basal or non-Basal, and CO or CF classification (d) Bar plots showing the average expression of all genes in individual basal networks per basal subtype group, in CO or CF cells. (e) Immunostaining for KRT5 (green) and PCNA (red) in sections from CF and CO lung tissue. Nuclei are stained with DAPI. Arrow indicate points of interest, while insets show magnification of the basal cell layer. (f) Quantification of KRT5+ PCNA+ basal cells in CO and CF. (g) Expression distributions of cell cycle genes in CO and CF cells, in the proliferating Basal2 subtype.

Using the gene network analysis approach, we defined 10 gene expression networks that were differentially regulated between CO and CF samples and were prominent in basal cells. Eight networks (Net B1-B4, B7-B10) were higher in CO samples and two networks (B5 and B6) were higher in CF samples (Fig 4b, Supp Fig 5, Supp Table4). The CF-enhanced B5 and B6 networks are related to surfactant metabolism and immune function, and, interestingly, were expressed in the smallest proportion of basal cells, specifically in the Basal5 subtype for B5 and the Basal4 subtype for B6 (Fig 4b, Supp Fig 5). Many networks showing down-regulation in CF compared to CO samples demonstrated gene ontologies related to metabolic processes and oxidative stress, cell division, epithelial cornification, immune functions, and response to wounding (Fig 4c, Supp Fig 5). Networks B1, B2 and B8 were more highly expressed in CO compared to CF samples (Fig 4c,d) and may signify patient specific wound healing related to intubation. Several other molecular pathways were also downregulated in the basal cells of CF samples compared to CO, including those related to response to oxidative stress, and ATP synthesis (Net B2, B4, B10, Fig 4c,d). Strikingly, networks B3 and B7 revealed widespread downregulation of genes related to cell cycle in CF samples across all basal subtypes but most notably among cells of the Basal2 (proliferating) cluster (Fig 4b,c,d), which may be related to the fact that the number of Basal2 cells is lower in CF than CO samples.

Our finding of reduced proliferative capacity among basal cells of CF compared to CO airways has important implications for the ability of endogenous stem/progenitor cells to maintain the specialized epithelial lining of CF airways. To confirm the depletion of dividing basal cells in intact CF mucosa that are inferred from single cell RNA-Seq data, we analyzed immunofluorescent co-staining for a proliferative marker (PCNA) and the basal marker, KRT5, in the same proximal airway samples used for transcriptomic analysis. We found that the PCNA-proliferative index of KRT5-immunoreactive cells in CF proximal airways was significantly reduced compared to comparable airway regions of CO tissue (Fig 4e,f, Supp Fig 6). Furthermore, analysis of cell-cycle transitional state signatures in transcriptomes of the proliferative Basal2 cell cluster confirmed a general reduction in all phases of the cell cycle among CF samples compared to their CO counterparts (Fig 4g). Taken together, the reduction in proliferation of the basal cells of the surface hBE cells has important implications for airway repair in CF and cellular gene targeting of long-lived stem/progenitor cells in CF.

## Discussion

Taken together, we report both novel identification of proximal airway epithelial basal, secretory, and ciliated molecular subtypes and insights about the transcriptional differences at the single cell level between airways from CF and control subjects. Histological reports have described basal cell hyperplasia in the CF airways^28,29^, but this was not corroborated in our study. We attribute this discrepancy to the increased sampling power and sample access associated with our study, in which we observed similar proportions of basal cells in airways of CO and CF lung, but find a notable decrease in the dividing basal subtype in CF airways.

Interestingly, it is the secretory cell subtype associated with mucosal immunity that is most highly upregulated in CF. Also, we report that secretory cells account for the largest fraction of *CFTR* transcript expression among all cell populations in both control and CF samples. In addition, the ciliated cells were found to have an increased number of transitioning precursors in the lungs of CF donors compared to controls suggesting that they may have more plasticity than their counterparts from CO donors. Lastly, upon examining the hBE cell ALI model system, we found that the diversity of cell subtypes in ALI cultures is different to that found in fresh tissue, presumably due to the effect of a uniform culture microenvironment.

By leveraging the analysis of 38 patient samples across a 3-institution consortium and assessing gene expression patterns that are common between datasets, we have generated molecular atlases of control and CF proximal airway epithelium. This molecular atlas was used to examine CF-lung disease dependent changes in the transcriptional phenotype of lung epithelial cells but can also be utilized as a hypothesis generating tool for other airway conditions. Our data suggest that specific subtypes of the main airway cell types have potential to play a role in CF lung disease, although *in vitro* and *in vivo* validation is still needed to assess the functional potential of these cell subtypes in health and disease. These studies provide valuable, novel insights into the molecular pathogenesis of CF lung disease and have potential to impact development of new therapies to ameliorate CF-related airway dysfunction.

## Supporting information

Supplemental files combined

## Acknowledgements

We would like to thank Susan Reynolds for her helpful input in reviewing this manuscript. This work was supported by the Cystic Fibrosis Foundation (CFF) (GOMPER17XX0 (BNG), STRIPP15XX0 (BRS), CARRAR19G0 (GC), BOUCHE15R0 (SHR)), the Tobacco-related disease research program (TRDRP) (HIPRA 29IP-0597)(BNG), NHLBI (PO1 HL108793)(BRS), NIH grant DK065988 (SHR) and a grant from Celgene/BMS (BRS). J. L. was supported by the UCLA Tumor Cell Biology Training Program (USHHS Ruth L. Kirschstein Institutional National Research Service Award # T32 CA009056); S. S. by the UCLA Broad Stem Cell Research Center – Rose Hills Foundation Training Award and currently the UCLA Dissertation Year Fellowship. KP and BNG were supported by the UCLA Broad Stem Cell Research Center, the David Geffen School of Medicine, and the Jonsson Comprehensive Cancer Center, and KP by NIH (P01 GM099134). The research of KP was also supported in part by a Faculty Scholar grant from the Howard Hughes Medical Institute. CJA and DWS were supported by UCLA Medical Scientist Training Program grant (NIH NIGMS GM008042), the NIH/NCI NRSA Predoctoral F31 Diversity Fellowship F31CA239655 (CJA), the UCLA Eli & Edythe Broad Center of Regenerative Medicine and Stem Cell Research Training Grant (CJA), the T32 National Research Service Award in Tumor Cell Biology CA009056 (CJA),

## Author Contributions

**GC, JL, JM** designed and performed experiments, analyzed the data and prepared the manuscript

**SS, ZL, AP, BK, CJA, BAC, PV, CY, DWS, EI, TMR, EW, AS, MM** assisted in tissue handling, sampling, processing, and sorting for single cell RNA seq

**SHR, EKV, ALR, MM** provided expertise and/or tissue analysis

**KP, JM, BRS, BNG** supervised the study and prepared the manuscript

All authors reviewed and edited the final manuscript.

