## Supplemental files combined for "A molecular atlas of proximal airway identifies subsets of known airway cell types revealing details of the unique molecular pathogenesis of Cystic Fibrosis"

**Supplemental Figure Legends**

**Supp Fig 1**

(a) Visualization of the distribution of cells from the three institutions in the integrated embedding, showed by institution and (b) by samples of origin, visualized by UMAP. (c-f) Network distributions with differences between institutions, visualized by UMAP. (g) Major cell types identified using previously described markers, visualized by UMAP. (h) Ionocyte and NE cell clusters analyzed independently of other cell types, visualized by UMAP. (i) CO and CF sample contribution to cell populations and subclusters, visualized by a stacked column chart. (j) Signatures of major cell types in ALI cells, created using previously published ALI gene lists, shown by violin plots. (k) Distribution of major cell type proportions in freshly isolated and ALI datasets. (l) Proportion of *CFTR* expressing cells, key at right, visualized by a stacked column chart.

**Supp Fig 2**

(a) For each cell, the average mean expression of the genes in a given network is shown, visualized on a UMAP. Cells are split by Secretory or non-Secretory, and CO or CF classification

(b) Bar plots showing the average expression of all genes in individual secretory networks per secretory subtype group, in CO or CF cells.

**Supp Fig 3**

(a) For each cell, the average mean expression of the genes in a given network is shown, visualized on a UMAP. Cells are split by Ciliated or non-Ciliated, and CO or CF classification

(b) Bar plots showing the average expression of all genes in individual ciliated networks per ciliated subtype group, in CO or CF cells.

**Supp Fig 4**

(a-j) For distinct categories of genes related to cilia biogenesis, the expansion of cilia gene expression is shown by violin plots and UMAP, indicating the changes in CO and CF for each cell subtype.

**Supp Fig 5**

(a) For each cell, the average mean expression of the genes in a given network is shown, visualized on a UMAP. Cells are split by Basal or non-Basal, and CO or CF classification

(b) Bar plots showing the average expression of all genes in individual basal networks per basal subtype group, in CO or CF cells.

**Supp Fig 6**

(a) Representative IF images of airways showing KRT5 (green) and PCNA (cyan), all nuclei are counterstained with DAPI (blue) in the merged image. (b) Representative examples of watershed segmentation for isolated KRT5 and PCNA staining. (c) Representative images indicating counting of KRT5 (green) and PCNA (cyan) expressing cells in the segmented images. Red and yellow boxes highlight areas that provide 4x zoomed images. (d) Segmentation data assumes a normal distribution. Each data point represents a possible cell and its corresponding area. Red line represents the mean area of the data and black line represents two standard deviations above the mean area. Representative tiles scan regions taken at 20x magnification for non-CF (e) and CF (f) subjects stained for KRT5 (green), PCNA (cyan) and nuclei are counterstained with DAPI (blue). Dimensions of the airways are indicated by the white lines.

**Supp Fig 7**

Representative FACS for isolation of epithelial cells to use in scRNAseq with 10X Genomics. Cell debris were excluded on the basis of FSC-A versus SSC-A, then doublets were removed using Trigger Pulse Width versus FSC-A (Influx). Dead cells were identified and excluded on the basis of staining with DAPI. Negative gating for CD45, CD31, and CD235a, combined with positive gating for EPCAM (CD326) were used to identify epithelial cells.

Supplemental Fig. 1

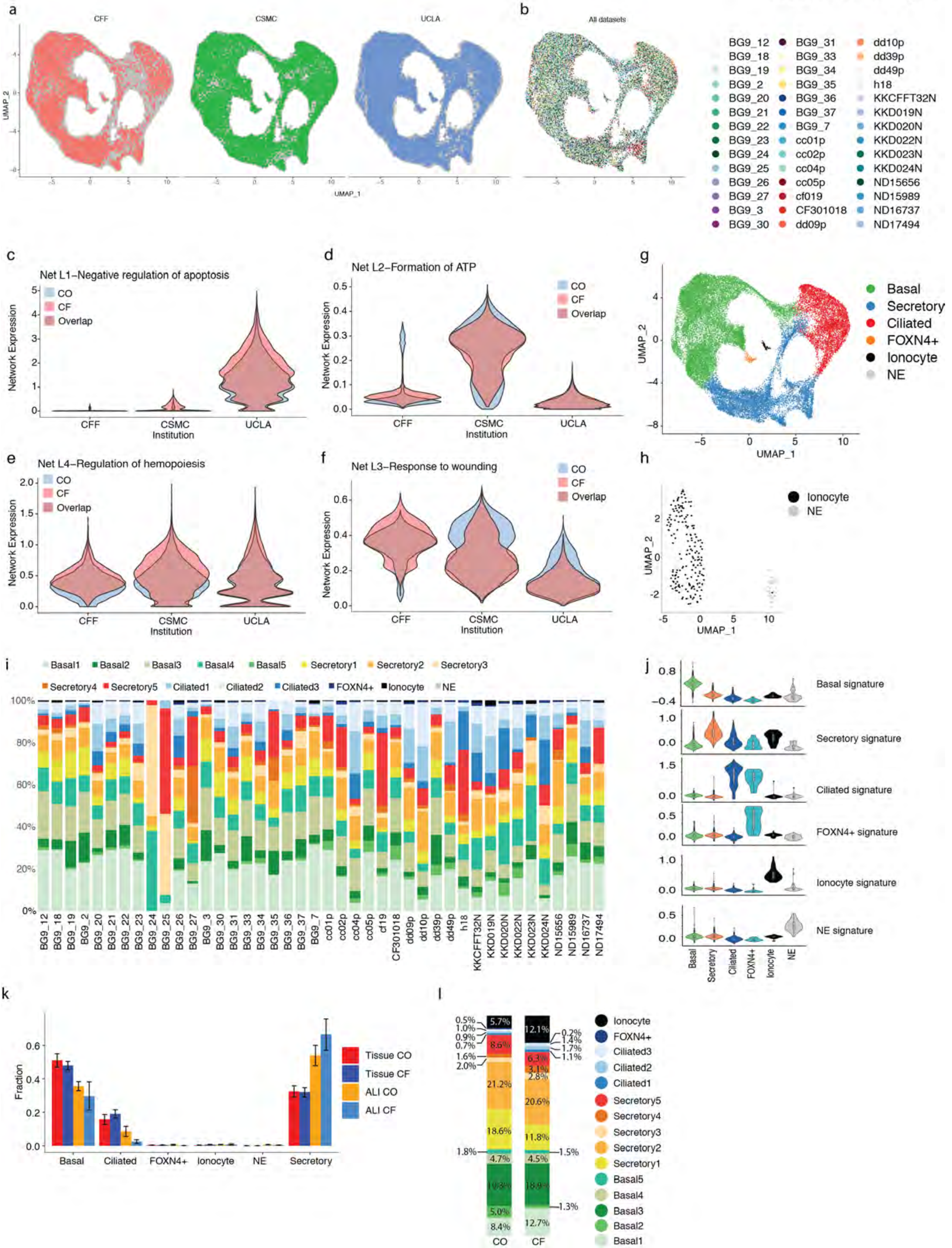

Supplemental Fig 2

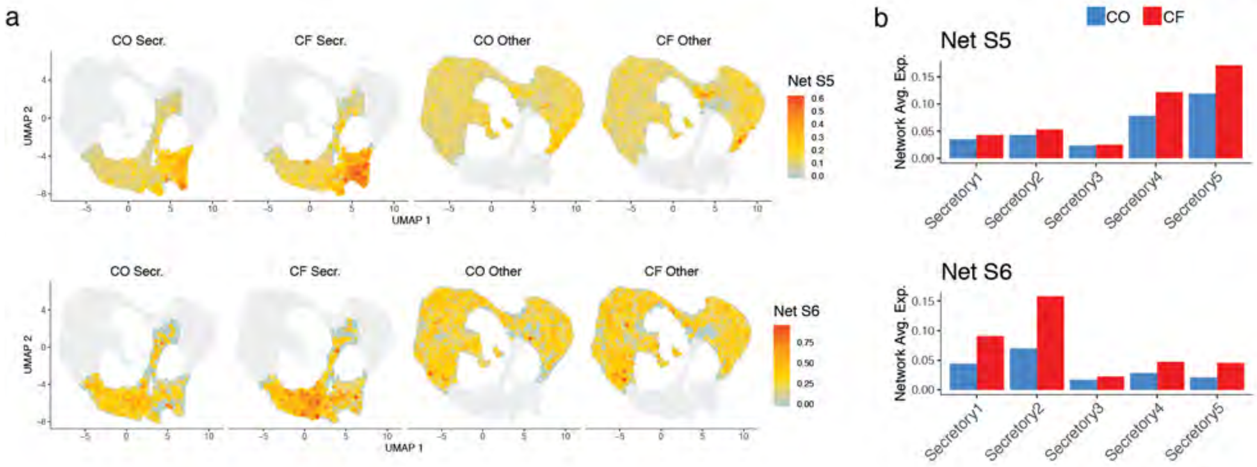

Supplemental Fig 3

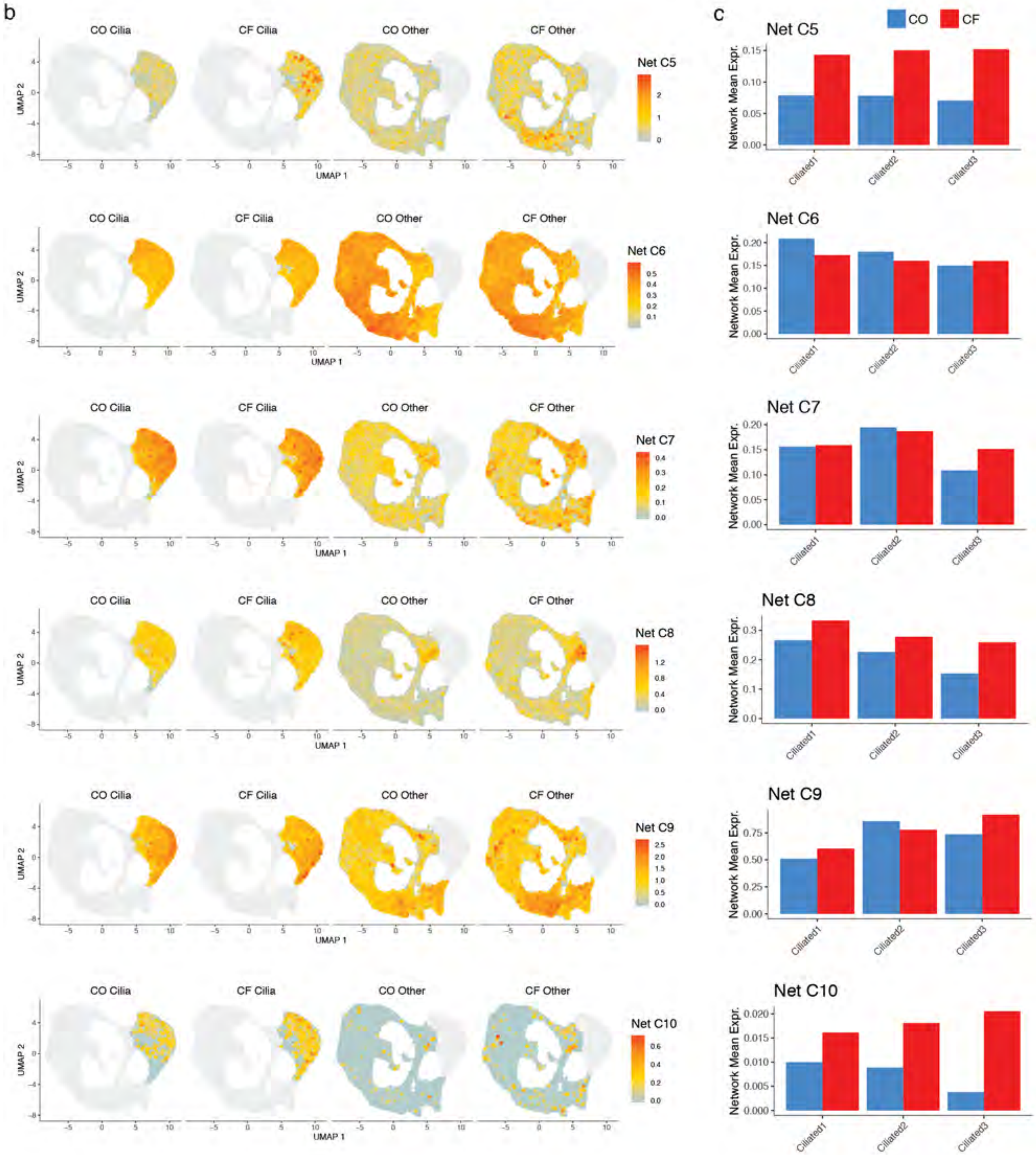

Supplemental Fig. 4

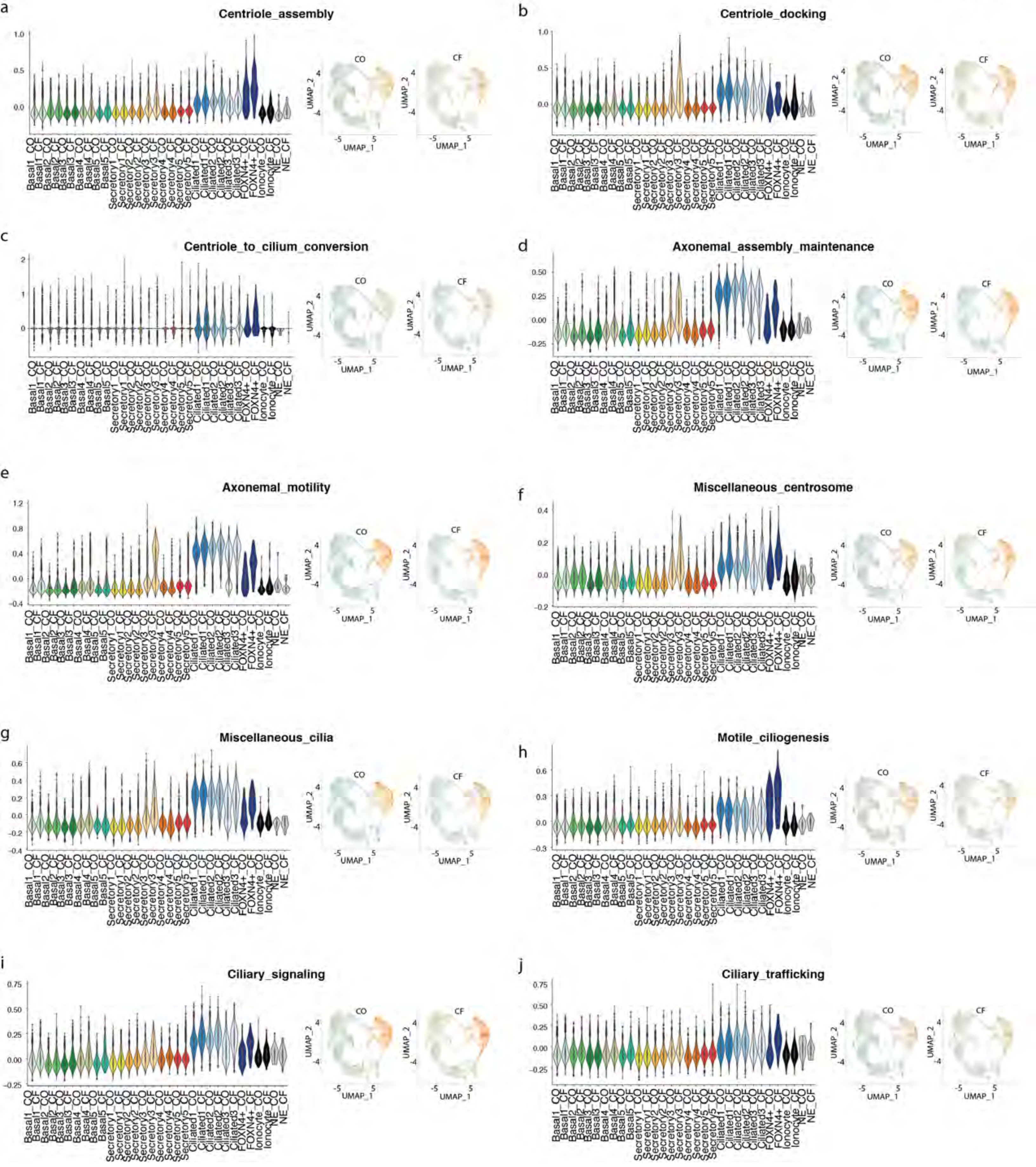

Supplemental Fig 5

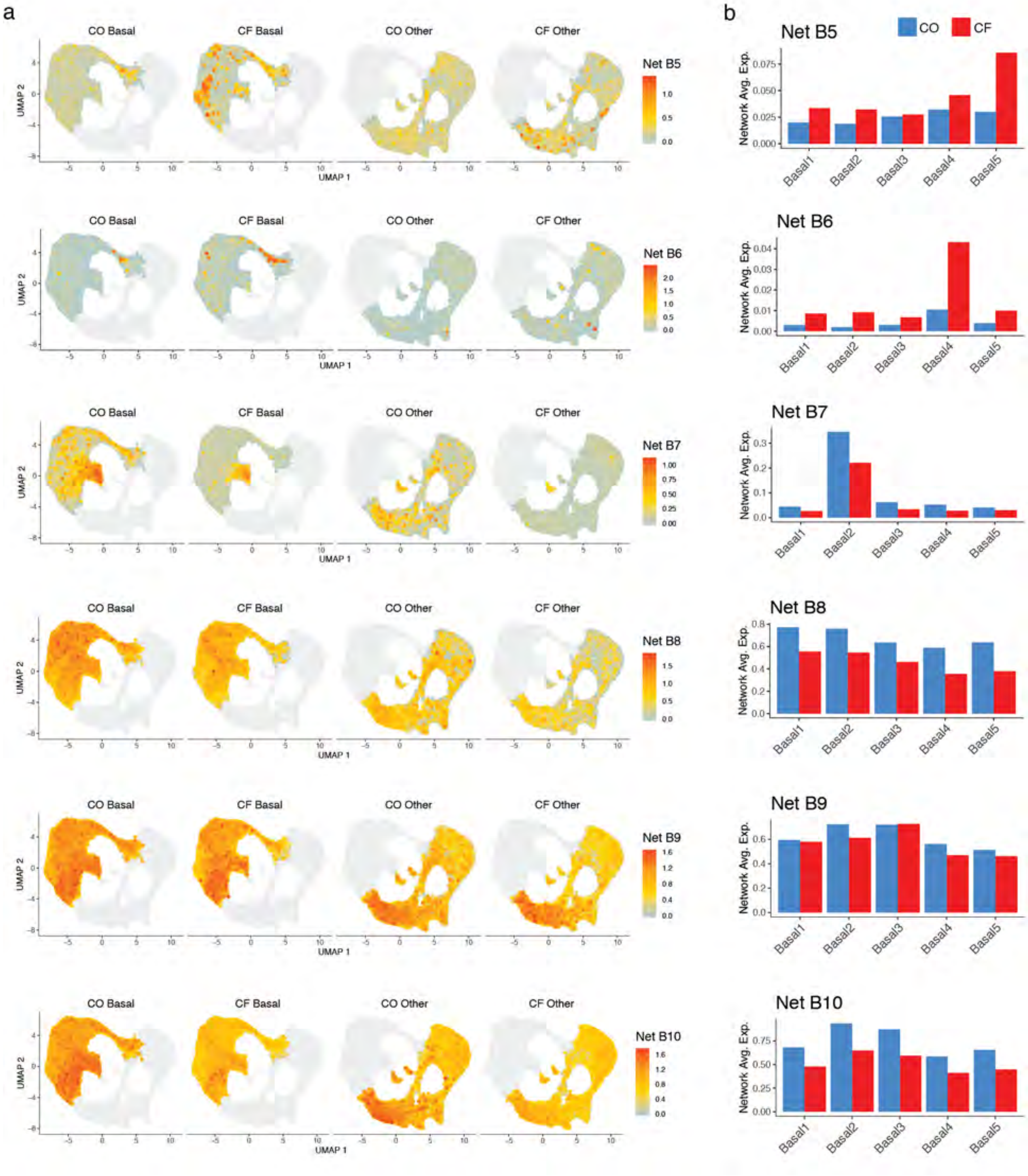

Supplemental Fig. 6

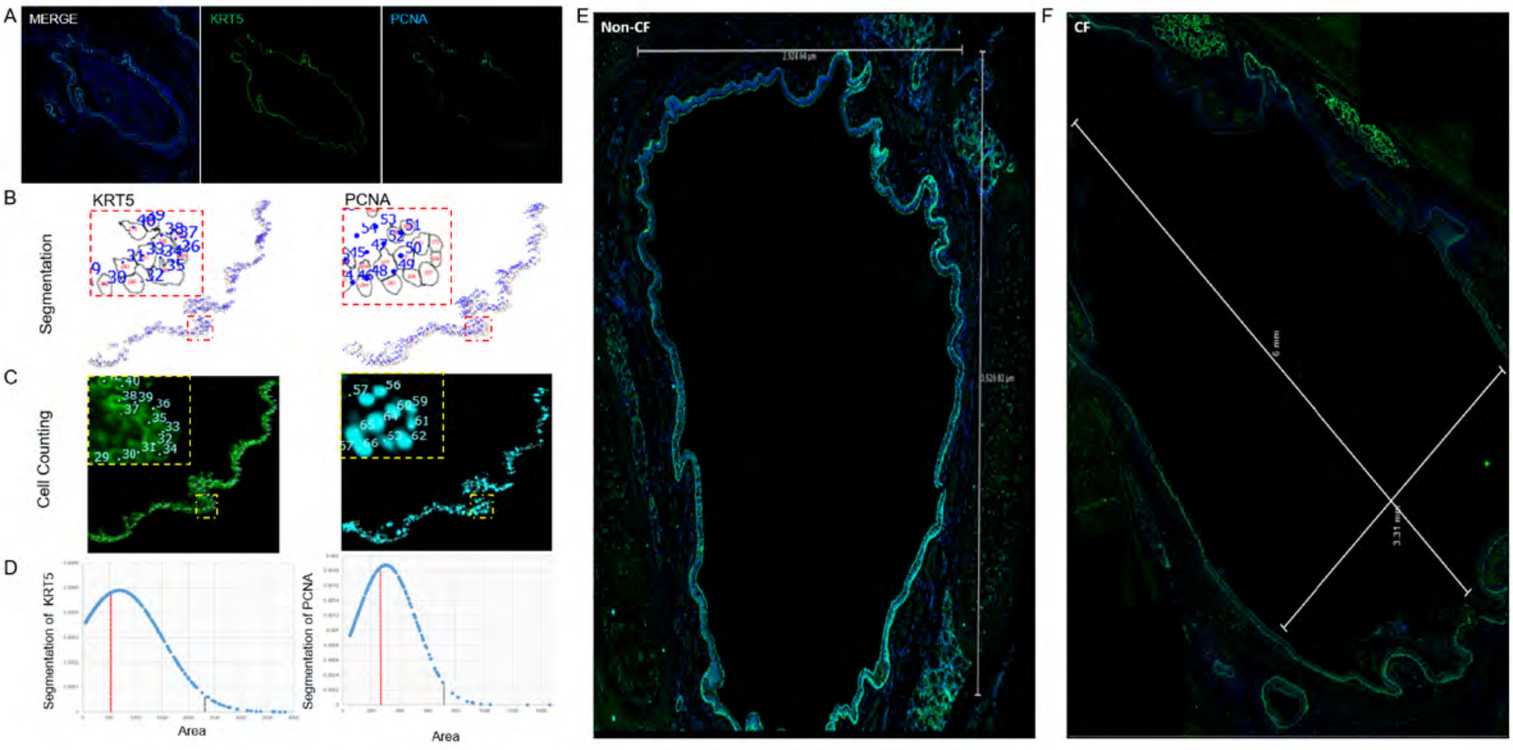

Supplemental Fig. 7

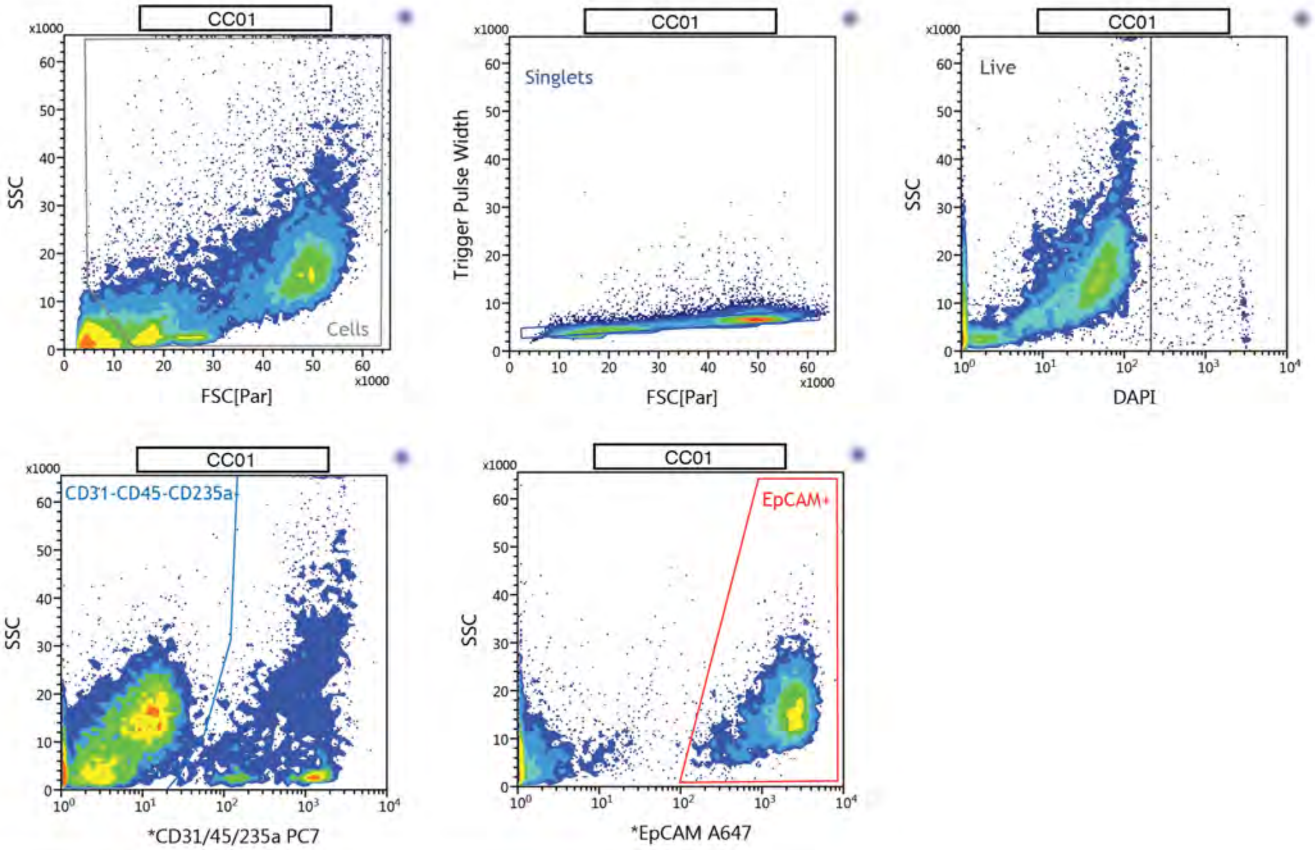


Populations: CC01

| Populations | Events | % Total | % Parent |
|---|---|---|---|
| All Events | 17,448 | 100.00% | #### |
| Cells | 12,827 | 73.52% | 73.52% |
| Singlets | 10,508 | 60.22% | 81.92% |
| Live | 10,269 | 58.85% | 97.73% |
| CD31-CD45-CD235a- | 7,494 | 42.95% | 72.98% |
| EpCAM+ | 4,306 | 24.68% | 57.46% |
